# Hydrocarbon effects on cell survival and antioxidant system of breast tumorigenic and non-tumorigenic cells

**DOI:** 10.1101/2024.11.08.622500

**Authors:** Mariana Mardirosian, Marianela Lasagna, Mariel Núñez, Tamara Galarza, Nuria Espert, Andrés Venturino, Claudia Cocca

## Abstract

Over the last decades, environmental pollution with polycyclic aromatic hydrocarbons (PAH) has risen due to human development and industrial activities. The consequences of potential contamination with oil or any of its components must be evaluated. Here, we faced the effects on cell growth and detoxification that might arise from the exposure to hydrocarbons sourced from contaminated waters. MCF-7 and MDA-MB-231 human breast cancer cells and/or MCF-10A epithelial non-tumorigenic cells were exposed to water-accommodated oil fraction (WAF) or anthracene as a PAH currently found in the environment to evaluate clonogenicity, viability, GST and CAT activities. WAF decreased MDA-MB-231 cells clonogenicity and MDA-MB-231 and MCF-7 cells viability. Anthracene significantly reduced non-tumorigenic cells’ viability and clonogenicity, without affecting the survival of tumorigenic cells. Both tumorigenic cells responded mainly by activating the antioxidant system through the increment of CAT when exposed to low concentrations of WAF. In both cell lines, WAF also increased GST. Anthracene exposure significantly decreased CAT activity of the three cell lines evaluated. GST activity decreased 38% after 28 µM anthracene exposure in MDA-MB-231 cells (p<0.05) and 15, 17 and 21% after 7 (p<0.05), 14 and 28 µM (p<0.01) in MCF-10A cells, while in MCF-7 cells 28 µM anthracene increased GST activity by 75% (p<0.001). We conclude that the sensitivity of cells to the different contaminants evaluated is affected by the degree of cellular tumorigenicity and exposure to WAF or anthracene may exacerbate malignant conditions. The integration of multiple parameters, analyzed through multivariate methods, offers a comprehensive evaluation of toxicant effects.

**HIGHLIGHTS:**

- WAF and anthracene affect tumorigenic mammary cells viability and clonogenicity
- Non tumorigenic mammary cells show more sensitivity to anthracene exposure
- WAF activates antioxidant and detoxicant system in tumorigenic mammary cells
- Anthracene decreased CAT activity of tumorigenic and non-tumorigenic mammary cells
- Anthracene affected GST activity of tumorigenic and non-tumorigenic mammary cells

## INTRODUCTION

The development of the oil industry and the growing demand for its products worldwide have increased the vulnerability of the environment to the harmful effects of oil pollution. Although the extractive process is controlled, incidents of spills and contamination of soil and water may occur. Several studies analyze the effects of water contamination with crude oil by using a water-accommodated fraction (WAF), which is the fraction mainly responsible for the toxic effects after spill dissipation [1]. WAF consists of a complex mixture of hydrocarbons including several polycyclic aromatic hydrocarbons (PAH), such as naphthalene, fluoranthene, pyrene, anthracene, phenanthrene, among others, depending on the petroleum extraction area [1–5]. Exposure of human and animals to PAH can trigger a variety of cellular responses, including, apoptosis, alterations in antioxidant metabolism, oxidative stress, chromosomal damage, genotoxicity, carcinogenicity, reproductive and endocrine disrupting effects, immunotoxicity, neurotoxicity, teratogenicity and cytotoxicity, among other alterations [6–8]. The type and severity of responses generated depend on a variety of factors [9] like the relative toxicity of the chemical, duration of exposure, and health status, among other factors [10]. For example, Drwal et al. (2017) observed an action dependent on the cell type after PAH exposure: a pro-apoptotic effect (increased Bax and caspase-3) in human placental BeWo cells and an anti-apoptotic effect (decreased Bax and increased cdk2 and cyclin D1) in human placental JEG-3 cells. Also, naphthalene, pyrene, and phenanthrene exhibited an endocrine disrupting effect in JEG-3 cells, but not in BeWo cells [11]. It has also been reported that skin, lungs, pancreas, esophagus, bladder, colon, and female breast are predisposed to tumor development due to long-term exposure to PAH [12]. Besides, preclinical studies have found a relationship between PAH exposure, oxidative stress and atherosclerosis [13,14]. However, there is very little information available on individual PAH; most of the information is for the entire PAH group [15]. Some PAH, such as anthracene, can be found in multiple environmental matrices. Anthracene is a PAH considered potentially harmful, whose environmental monitoring is considered essential. Based on the environmental risks and extent of use of PAH, the USEPA included anthracene among the 16 PAH considered as priority pollutants [16]. Anthracene is a highly hydrophobic molecule, with great chemical stability and insufficient biodegradability [17].

Due to their lipophilic nature, PAH, pesticides and other organic compounds can accumulate in cell membranes, tissues with high fat content and other cellular lipid deposits enhancing their bioavailability in organisms [12]. For example, naphthalene has been shown to accumulate in the adipose tissue [18]. Ramos Nieto et al. reported a dose-dependent accumulation of chlorpyrifos in rat abdominal adipose tissue [19] and anthracene was found at high levels in the liver and human fat [20]. Human breast is a tissue composed of a stroma with high lipid content and epithelial and myoepithelial cells that compose its characteristic structure, therefore lipophilic contaminants like PAH can bioaccumulate and generate local effects. It is also known that PAH can alter oxidative metabolism and redox balance in normal and malignant tissues [9,12,21,22]. It is estimated that between 70% and 90% of all cancers are related to exposure to environmental risk factors [23]. Besides, breast cancer is one of the most common cancers among women, with an estimated global prevalence of 2.3 million by 2030 [24]. However, individual toxic chemicals have not yet been identified as risk factors for breast cancer. On the one hand, because we are exposed to a complex environment that changes with the season of the year. On the other hand, breast cancer risks include not only environmental factors but also the age when the exposure occurs and the properties of the pollutants. Furthermore, Balise et al. (2016) reviewed available evidence indicating that numerous chemicals generated in oil and gas operations have the potential to modulate estrogen receptor (ER). Predominantly, studies employing reporter gene assays in human cell lines, complemented by affinity assays in human cells, have demonstrated that various hydrocarbons associated with oil and gas activities can directly interact with the ER. Given that several studies documented disruption of the ER via either agonistic or antagonistic mechanisms, the cumulative evidence supports the conclusion that oil and gas hydrocarbons contribute to the introduction of endocrine-disrupting substances into the environment [25].

Given the ubiquity and toxicity of PAH, we are particularly interested in evaluating the potential effects of polycyclic compounds on those cells that may be surrounded and influenced by a lipid microenvironment with a high probability of bioconcentration of these contaminants, with harmful consequences. Therefore, in this study, we focused on elucidating how WAF or anthracene, a PAH selected as a model hydrocarbon due to its ubiquity, can affect parameters associated with redox imbalance that could have implications on breast cells derived from normal or tumor tissue, with different ER_ status, to determine if these could be factors that alter survival and/or transformation.

## MATERIALS AND METHODS

### 1. WAF preparation from crude oil and hydrocarbon determinations

WAF was prepared from crude oil extracted from Chachahuen field, located in the Neuquén basin at North Patagonia (37°19’55.3’S 68°56’03.5’’W), following the methods outlined by del Brio et al. [4]. WAF hydrocarbon analyses were performed as previously described [4] and according to the Method for the Determination of Extractable Petroleum Hydrocarbons of the Massachusetts Department of Environmental Protection (MADEP), EPH–04 Revision 1 (MADEP - Massachusetts Department of Environmental Protection-State Agency, 2004).

### 2. Cell culture and treatment

Human breast cancer cells MCF-7 (HTB-22, ATCC) and MDA-MB-231 (HTB-26, ATCC) and non-tumorigenic epithelial cells MCF-10A (CRL-10317, ATCC) were cultured as previously described [26]. All cultures were kept at 37 °C with 5% CO_ and a humidified atmosphere. The cells used in all experiments were between passage 8 and 20 after thawing from the original stocks. WAF was filtered with a 0.2 µm nitrocellulose filter and mixed with cell growth medium to prepare serial dilutions from the original solution (1/500, 1/250, 1/100, 1/50 and 1/25 that corresponded to 7.32, 14.64, 36.6, 73.2 and 143.4 µg/L total hydrocarbon concentrations). Anthracene was dissolved in ethanol and 3.5, 7, 14 and 28 µM solutions were prepared in the growth media. In concentration-response experiments with anthracene, control cells were treated with the maximal concentration of ethanol used in the anthracene-exposed cells, which never exceeded 0.5% (v/v). Ethanol did not affect any of the parameters evaluated (data not shown).

### 3. Clonogenic assay

MCF-10A (3 x 10^3^ cells/well), MCF-7 (3 x 10^3^ cells/well) and MDA-MB-231 (1.5 x 10^3^ cells/well) were seeded in 6-well plates and incubated for 24 h for cell attachment. Cells were then treated for 7 days with either WAF (dilutions from 1/500 to 1/25 in culture medium) or anthracene (solutions from 3.5 to 28 µM), fixed with ice-cold methanol for 20 min, and stained with toluidine blue in 10% (v/v) aqueous ethanol. The clonogenicity was evaluated by counting colonies having at least 50 cells and the value was expressed as a percentage of the total number of colonies obtained in presence of the vehicle alone.

### 4. Cell viability

MCF-10A (1.5 x 10^3^ cells/well), MCF-7 (1.5 x 10^3^ cells/well) and MDA-MB-231 (1.0 x 10^3^ cells/well) cells were seeded in 96-well plates and incubated for 24 h to enable cell attachment. Cells were then exposed to WAF (dilutions from 1/500 to 1/25 in culture medium) or anthracene (from 3.5 to 28 µM) for 72 h. Cell viability was evaluated with 3-(4,5-dimethylthiazol-2-yl)-2,5-diphenyltetrazolium bromide (MTT) according to the manufacturer’s instructions. Briefly, cells were incubated with 0.5 mg/mL of MTT for 2 h at 37 °C. The resulting formazan crystals were dissolved in 100 µL of DMSO, and the absorbance was measured at 570 nm using a microplate reader. The absorbance value in each well was expressed as a percentage of the values obtained in control cells.

### 5. *Catalase and* glutathione S-transferase *activity*

Cells were exposed to WAF (dilutions from 1/500 to 1/25 in culture medium) or anthracene (from 3.5 to 28 µM) for 72 h, trypsinized and suspended in a phosphate buffer (KH_2_PO_4_/K2HPO4 50 mM, pH 7.8), followed by ultrasonic disruption and centrifugation at 10,000 x g for 10 min at 4 °C.

5.1 Catalase (CAT) activity was measured spectrophotometrically by monitoring the disappearance of H_2_O_2_ at 240 nm in an UV/visible spectrophotometer at 25_°C (Shimadzu, Kyoto, Japan) as we previously reported [27] with slight modifications. Briefly, the reaction mixture contained 50 mM sodium phosphate buffer (pH 7), 25 mM H_2_O_2_ and 50 μL of supernatant, in a total volume of 1 mL, and the absorbance was recorded for 1_min. The reaction mixture absorbance was controlled to be 1 at 240_nm. Specific activity was expressed as International Units (IU) of CAT per mg of protein, using a molar extinction coefficient of 40 M^−1^_cm^−1^. Protein concentration was determined by Bradford assay [28]. Results were expressed as a percentage of the specific activity in the control group.

5.2 Glutathione S-transferase (GST) activity was evaluated as previously described [27,29]. The enzymatic reaction was started adding 10_μL of the supernatant to the reaction mixture. Absorbance was recorded continuously at 340_nm for 1_min in a UV/visible spectrophotometer at 25_°C (Shimadzu, Kyoto, Japan). Rate measurements were corrected for the non-enzymatic reaction. The activity was expressed as μmol of CDNB conjugated/min/mg of protein, using a molar extinction coefficient of 9.6 M^−1^_cm^−1^. Results were expressed as a percentage of the specific activity in the control group.

## 6. Statistical Analysis

6.1 For all the experiments, at least three independent replicates were used to determine the mean ± SEM. One-way ANOVA along with Dunnet’s *post-hoc* test was used to compare between exposed and control groups. The data were analyzed by the GraphPad Prism 7.0 software (GraphPad Software Inc., Philadelphia, PA, USA). All *p* values less than 0.05 were considered statistically significant.

### 6.2 Integrated analysis of the variability of the response to WAF and anthracene

#### 6.2.1 Principal Components Analysis

The response of cells to either WAF or anthracene at the cytotoxic and biochemical levels was analyzed using Principal Components Analysis (PCA) (NTSYS package). Mean values of all the effects evaluated to the whole range of WAF or anthracene concentrations were used for the multivariate analysis.

#### 6.2.2 Integrated Biomarker Response analysis

We calculated an integrated biomarker response (IBR) index corresponding to each WAF or anthracene concentration level using the ‘Integrated Biological Responses version 2’ approach [30]. The standardized and centered responses of the different mean values at each WAF or anthracene concentration were displayed in star plots to identify different response patterns, and the IBR indexes for WAF or anthracene were represented as a function of the effective concentration.

## RESULTS

### 1. Analysis of WAF composition

The composition of WAF extracted from Chachahuen crude oil is summarized in table 1. The concentration of total petroleum hydrocarbons (TPH) was 3.66 mg/L, mainly as C9– C36 n-alkanes, which comprised about 93% of hydrocarbon mass. Other compounds found were single-ring aromatic hydrocarbons (BTEX group: toluene, 0.17 mg/L) and PAH, particularly naphthalene and phenanthrene (0.011 and 0.005 mg/L respectively).

**Table 1:** WAF composition prepared from crude oil from the Chachahuen basin.

| Classification | Hydrocarbon | Concentration |
| --- | --- | --- |
| TPH | C9-C36 | 3.66 |
| BTEX | Toluene | 0.17 |
| PAH | Naphthalene<br>Phenanthrene | 0.011<br>0.005 |

### 2. Cell proliferation

No significant differences were observed in the clonogenicity of MCF-7 cells exposed to WAF compared to control cells. MDA-MB-231 cells significantly decreased their clonogenicity after exposure to 1/500, 1/50 and 1/25 WAF dilutions. No significant differences were observed in the clonogenicity of anthracene-treated MDA-MB-231 or MCF-7 cells compared to control. On the other hand, MCF-10A clonogenic capability significantly decreased 24, 29 and 32% after exposure to 7, 14 and 28 µM anthracene respectively (p<0.05 and p<0.01), in a dose-dependent manner (Figure 1).

**Figure 1:**
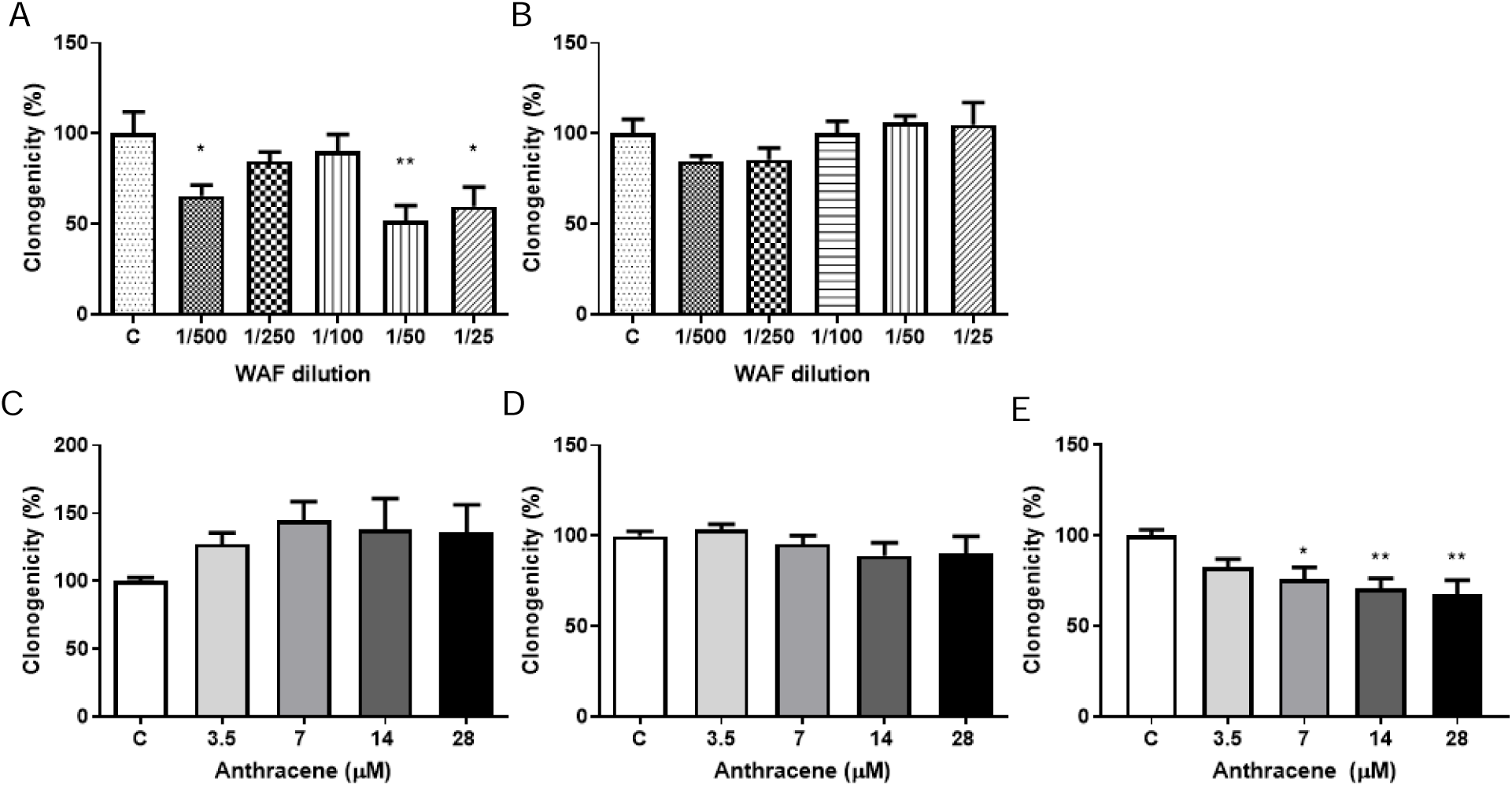
Effects of WAF or anthracene on clonogenic capability. MDA-MB-231 (A and C), MCF-7 (B and D) and/or MCF10-A (E) cells were treated for 7d with WAF dilutions (A and B) or anthracene (C, D, E) and the number of colonies with more than 50 cells was assessed. *p<0.05, **p<0.01. One-way ANOVA and Dunnet’s post hoc test. Results are expressed as a percentage of control. n= 3 independent experiments.

### 3. Cell viability

To evaluate the effect of WAF or anthracene on cell viability, cells were exposed for 72 h to WAF or anthracene, and MTT assay was performed. A significant decrease in cell viability was observed after the exposures to the lowest WAF solutions, of 18% in MCF-7 cells (1/500 and 1/250, p<0.01) and of 20% in MDA-MB-231 cells (1/500 and 1/100 dilutions, p<0.05), compared to control group (Figure 2 A-B). When MCF-7 and MDA-MB-231 cells were exposed to different concentrations of anthracene (3.5 to 28 µM) for 72 h, no significant differences in viability were observed compared to control. However, non-tumorigenic MCF-10A cells significantly decreased their viability by 9.4, 8.9 and 11% after exposure for 72 h to 7, 14 (p<0.01) and 28 µM anthracene (p<0.001) respectively, compared to non-exposed cells (Figure 2 C-D-E).

**Figure 2:**
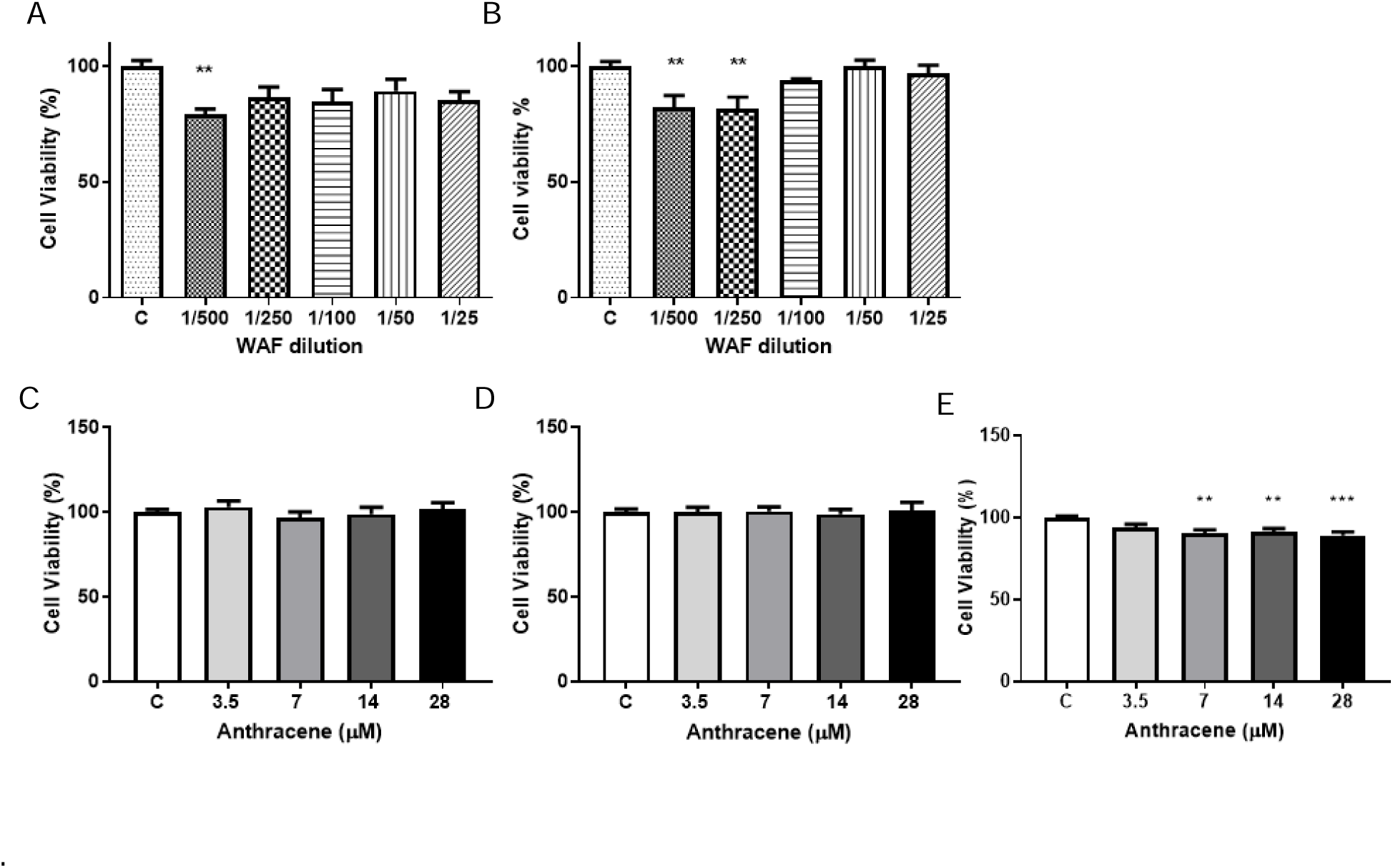
Effects of WAF or anthracene on cellular viability. MDA-MB-231 (A and C), MCF-7 (B and D) and/or MCF10-A (E) cells were treated for 72h with WAF dilutions (A and B) or anthracene (C, D, E) and MTT assay was performed. **p<0.01, ***p<0.001. One-way ANOVA and Dunnet’s post hoc test. Results are expressed as a percentage of control. n= 3 independent experiments.

### 4. CAT and GST activity

To assess the participation of the antioxidant and/or detoxifying systems after WAF (Figure 3) or anthracene (Figure 4) exposure, we studied the activities of CAT and GST enzymes. The most diluted WAF solutions significantly increased CAT activity of MCF-7 breast cancer cells from 100% (control group) to 170% (1/500) and 224% (1/250), after 72h of exposure (p<0.01 and p<0.001). Similarly, MDA-MB-231 cells exposed to 1/250 WAF significantly increased CAT activity by 45,9% compared to the control group (p<0.01). GST activity was also significantly increased for both cell lines after WAF exposure. In MCF-7 cells, GST activity increased by 27,9% after exposure to 1/250 WAF dilution (p<0.05), while for MDA-MB-231 cells, GST activity increased 42.3, 36.9 and 36.3% after 72h of exposure to 1/500, 1/50 and 1/25 WAF dilutions respectively, compared to control (p<0.05, Figure 3).

**Figure 3:**
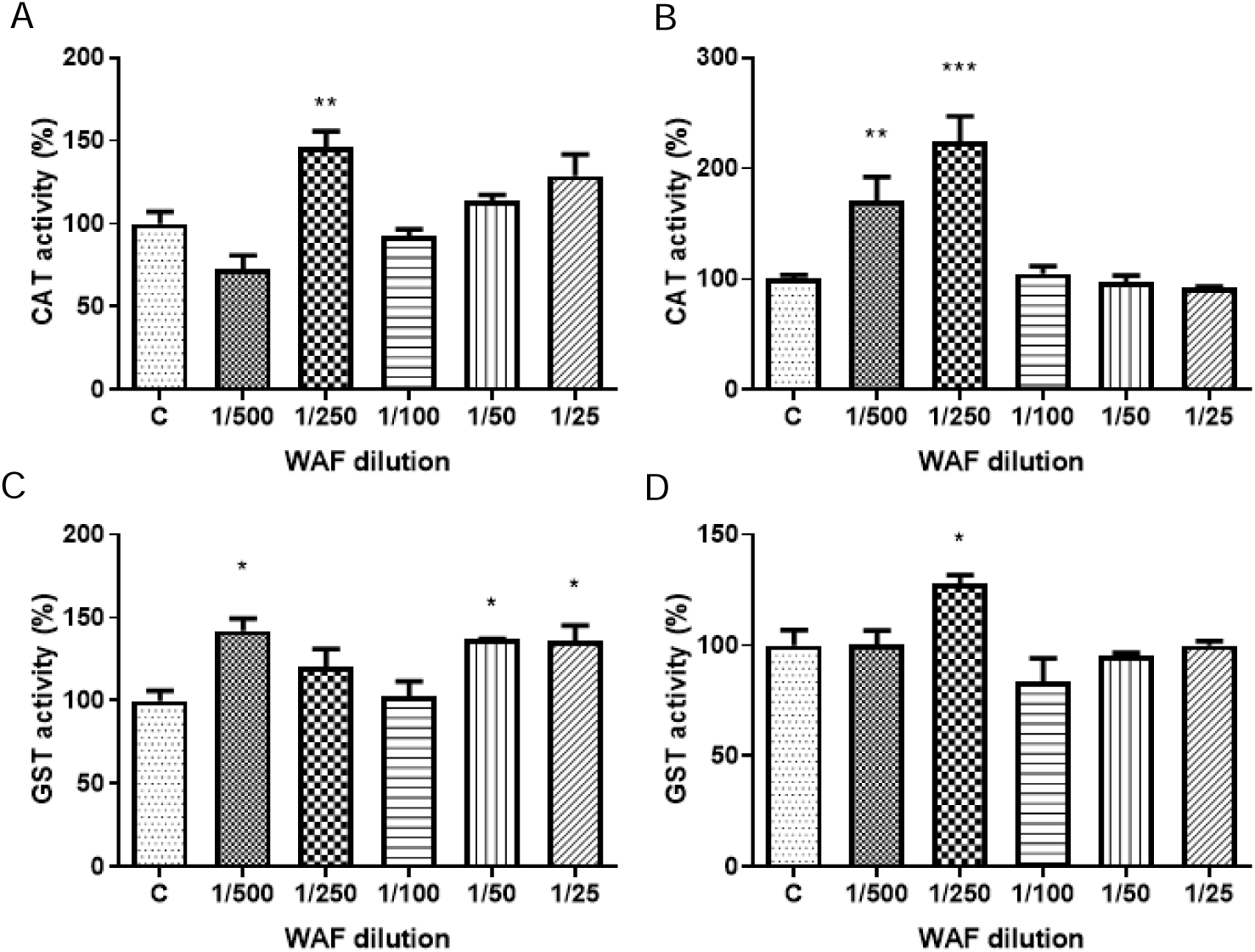
Effects of WAF on the antioxidant/detoxifying system. MDA-MB-231 (A and C) and MCF-7 (B and D) cells were treated for 72h with WAF dilutions to evaluate WAF effects on catalase (CAT) and glutathione S-transferase (GST) activities *p<0.05, **p<0.01, ***p<0.001. One-way ANOVA and Dunnet’s post hoc test. Results are expressed as a percentage of control. n= 3 independent experiments.

**Figure 4:**
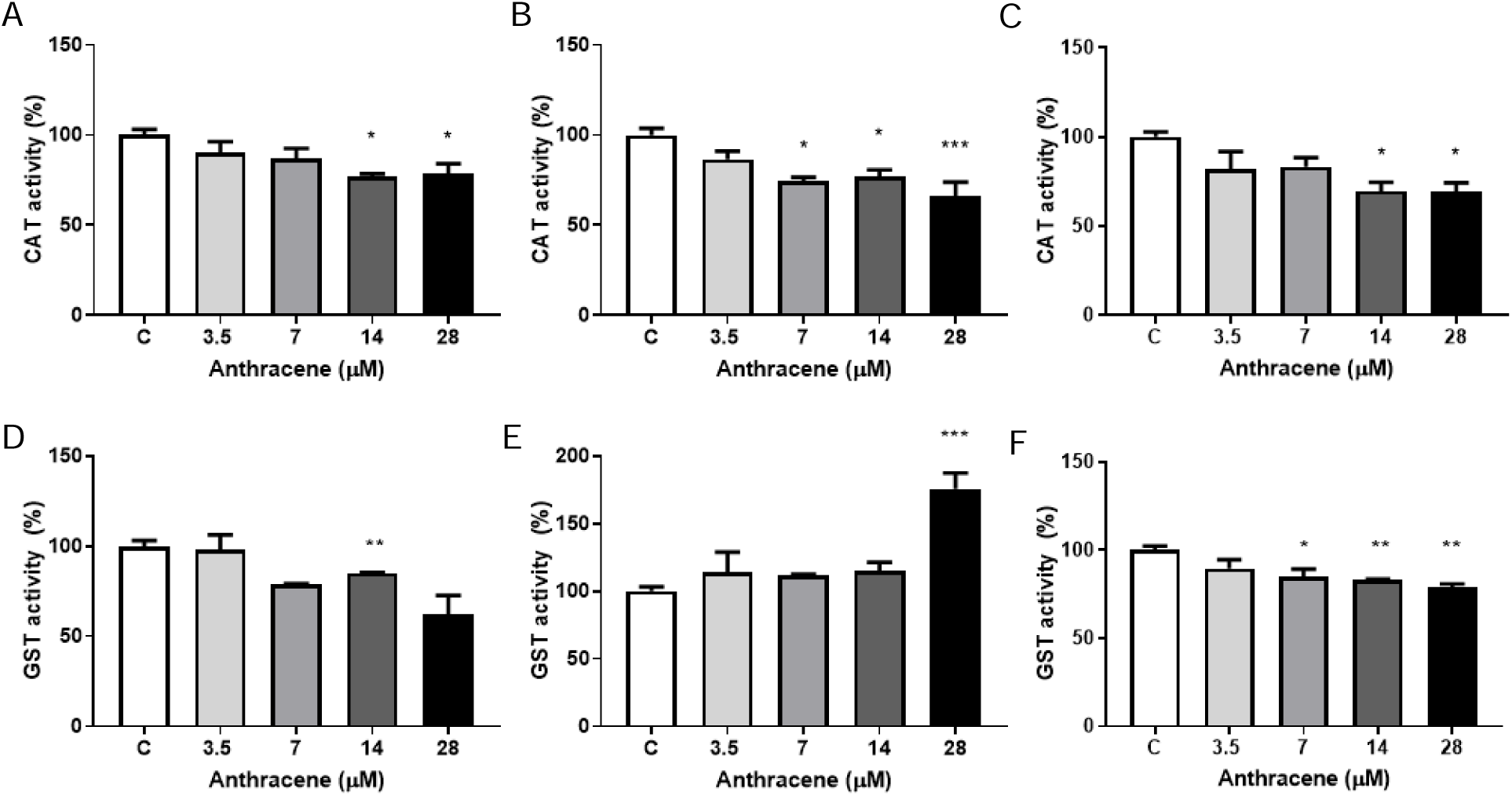
Effects of anthracene on the antioxidant/detoxifying system. MDA-MB-231 (A and D), MCF-7 (B and E) and MCF-10A (C and F) cells were treated for 72h with anthracene solutions (control, 3.5, 7, 14 or 28 μM) to evaluate anthracene effects on catalase (CAT) and glutathione S-transferase (GST) activities *p<0.05, **p<0.01, ***p<0.001. One-way ANOVA and Dunnet’s post hoc test. Results are expressed as a percentage of control. n= 3 independent experiments.

Figure 4 shows the effects of anthracene on CAT and GST activities. Exposure to the most concentrated anthracene solutions significantly decreased CAT activity of MCF-7, MDA-MB-231 and MCF-10A cell lines. In MDA-MB-231 cells CAT activity decreased 23 and 21% after exposure to 14 and 28 µM anthracene respectively, compared to control (p<0.05), while in MCF-7 cells exposure to 7, 14 and 28 µM decreased CAT activity by 26, 23 (p<0.05) and 34% respectively (p<0.001) and MCF-10A cells decreased CAT activity by 30.4 an 30.9% after exposure to 14 and 28 µM (p<0.05). GST activity decreased 38% after 28 µM anthracene exposure in MDA-MB-231 cells (p<0.05) and 15, 17 and 21% after 7 (p<0.05), 14 and 28 µM (p<0.01) anthracene exposure in MCF-10A cells, while in MCF-7 cells 28 µM anthracene increased GST activity by 75% (p<0.001).

### 5. Multivariate analyses of responses

We performed a multivariate statistical approach to assess the weight and possible correlations between cytological and biochemical responses, comparing the effects of WAF and anthracene in the different cell lines.

The Principal Components Analysis (PCA) performed on tumorigenic MCF-7 and MDA-MB-231 cells and non-tumorigenic MCF-10A cell lines were first analyzed together. This first analysis showed that in non-tumorigenic cells, all the related treatments were projected in the center, equidistant from all the response variables as equally affected by them all (see supplementary figure 1S). Then, only tumorigenic cells were analyzed, and both cell lines and contaminants clearly defined separate response patterns for WAF and anthracene (see supplementary figure 2S). Consequently, we performed separate PCA on the contaminants.

PCA on WAF exposures indicated that the first component C1 explained 64.5% of the total variability, with the contribution of the variables clonogenicity (CLON), Viability (Viab) to one side and GST to the opposite. In turn, C2 explained 24.8%, with the only contribution of CAT (Figure 5A). Projected treatments suggested that MCF-7 exposed to WAF was associated with high CLON or Viab values, while MDA-MB-231 was associated with high GST and opposite to CLON and Viab (in correlation with low values). Low WAF levels (1/500; 1/250) were close to high CAT activity for both cell lines.

**Figure 5:**
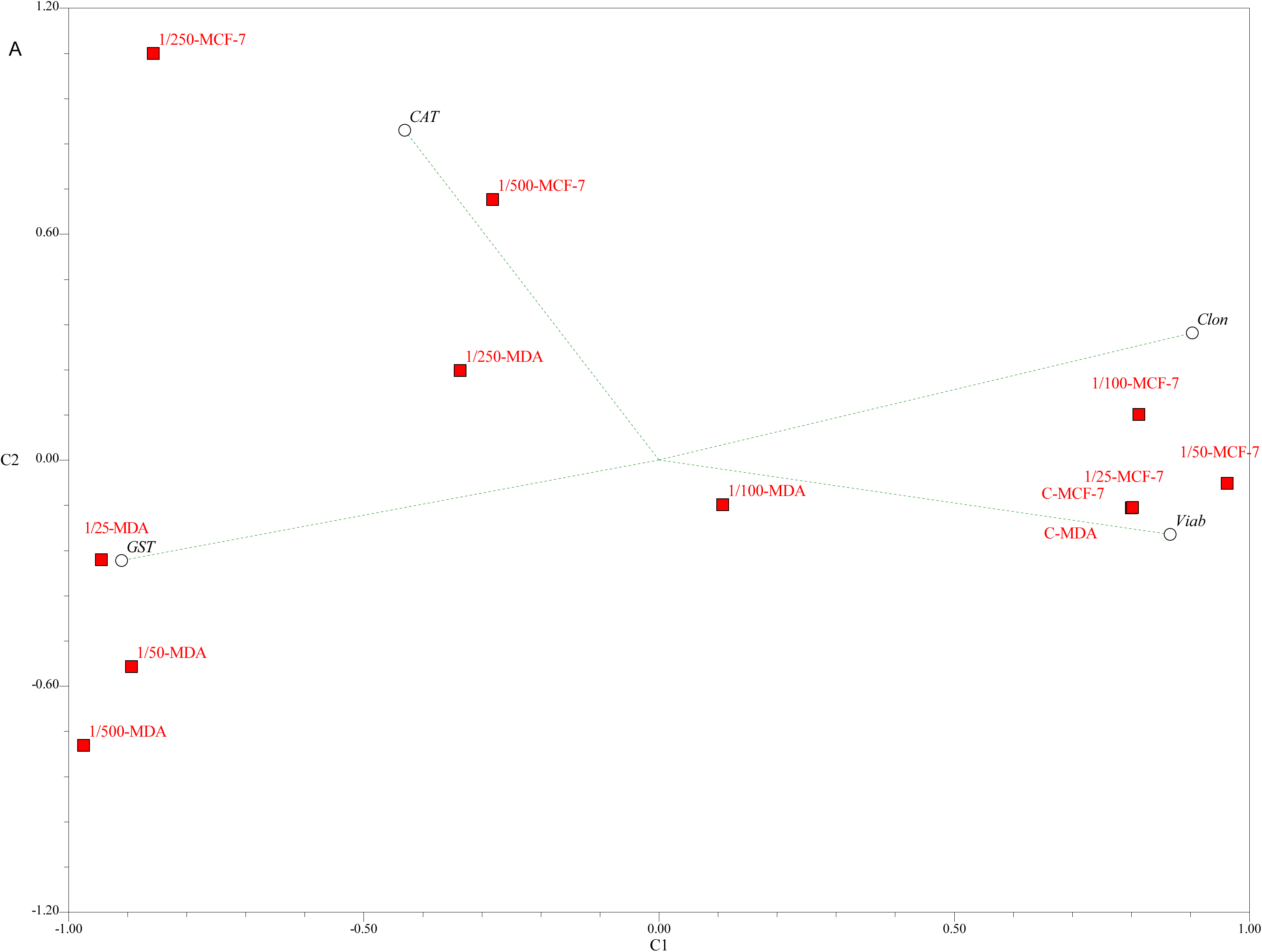
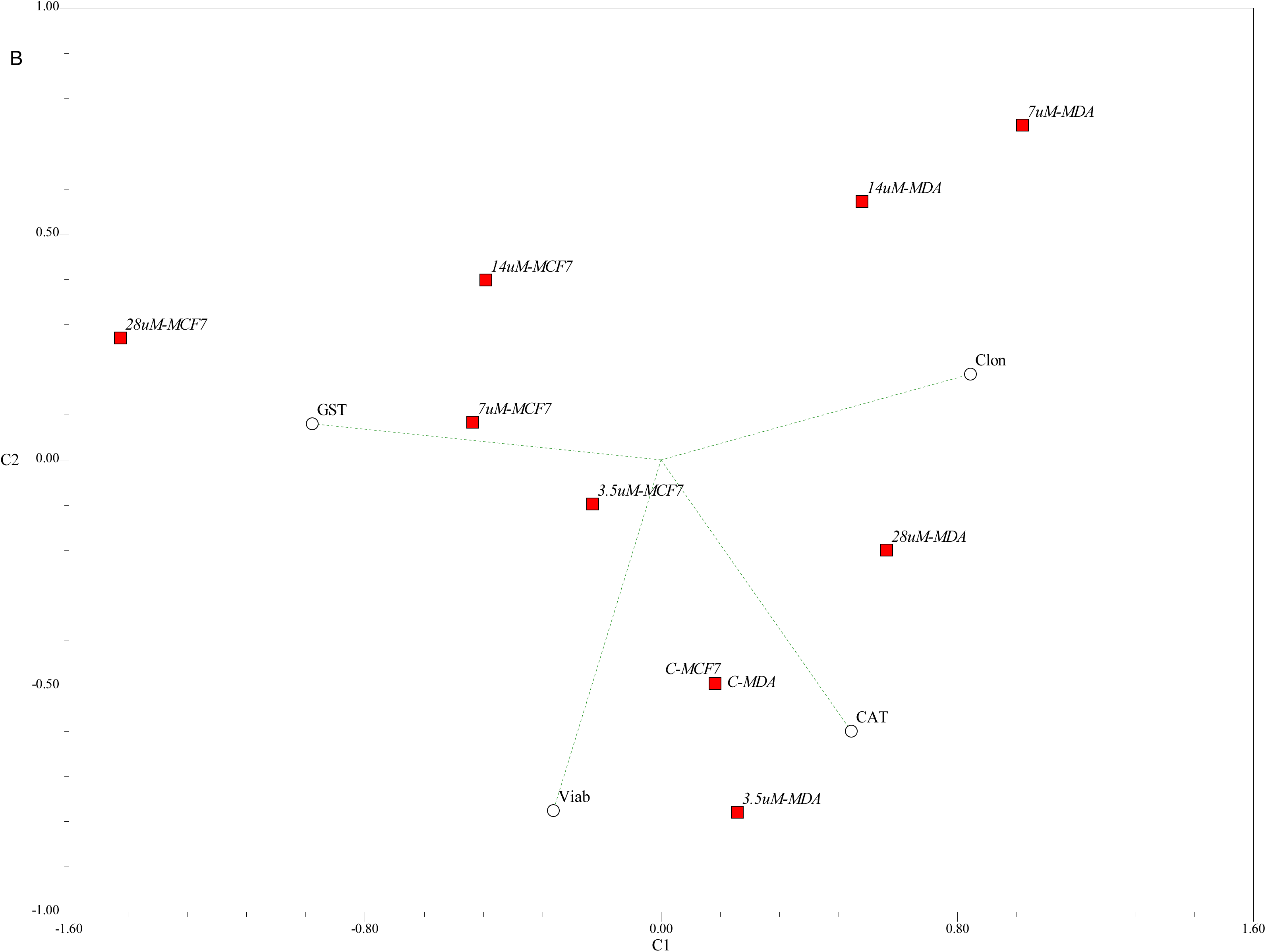
A - Principal Component Analysis on biomarker responses assessed in MDA-MB-231 and MCF-7 cells exposed to WAF; B - Principal Component Analysis on biomarker responses assessed in MDA-MB-231 and MCF-7 cells exposed to anthracene. CAT: catalase activity, GST: glutathione S-transferase activity, Viab: viability, Clon: clonogenicity. MDA: MDA-MB-231 cells, MCF-7: MCF-7 cells. WAF dilutions: 1/500, 1/250, 1/100, 1/50, 1/25. Anthracene concentrations: 3.5, 7, 14, 28 µM.

PCA on anthracene exposures resulted in a 48.4% of total variability explained by C1, with the contribution of CLON and GST to opposite sites, while C2 explained another 25.1% with the influence of Viab and CAT in the same direction (Figure 5B). When projected, exposed MCF-7 cells seemed more influenced towards high GST values, and exposed MDA-MB-231 cells seemed displaced to low GST and high clonogenicity and separated in C2 according to CAT activities.

Complementary, the Integrated Biomarker Response (IBR) approach displayed the differential behavior of the selected biomarkers with respect to WAF or anthracene exposure (Figure 6A-B). WAF caused an IBR with a biphasic dependence with the concentration, even different for MCF-7 or MDA-MB-231 cells. In turn, IBR was proportional to anthracene concentration in the range tested, and for the three cell lines. Star plots aided in understanding the variability of responses in the individual biomarkers along the different contaminant ranges. Low WAF levels influenced CAT and GST activity in MCF-7 cells increasing them, also causing lowered viability; in MDA-MB-231 cells, all WAF levels increased GST activity, and in general reduced clonogenicity and viability (Figure C-D). Anthracene proportionally reduced CAT activity in MCF-7 cells, and the highest level of the PAH increased GST activity; in MDA-MB-231 cells, it reduced CAT activity proportionally to the concentration, reduced GST activity at high levels and affected viability in a variable way. The effect of anthracene in MCF-10A cells was quite uniform, reducing all biomarker levels (Figure 6E-G).

**Figure 6:**
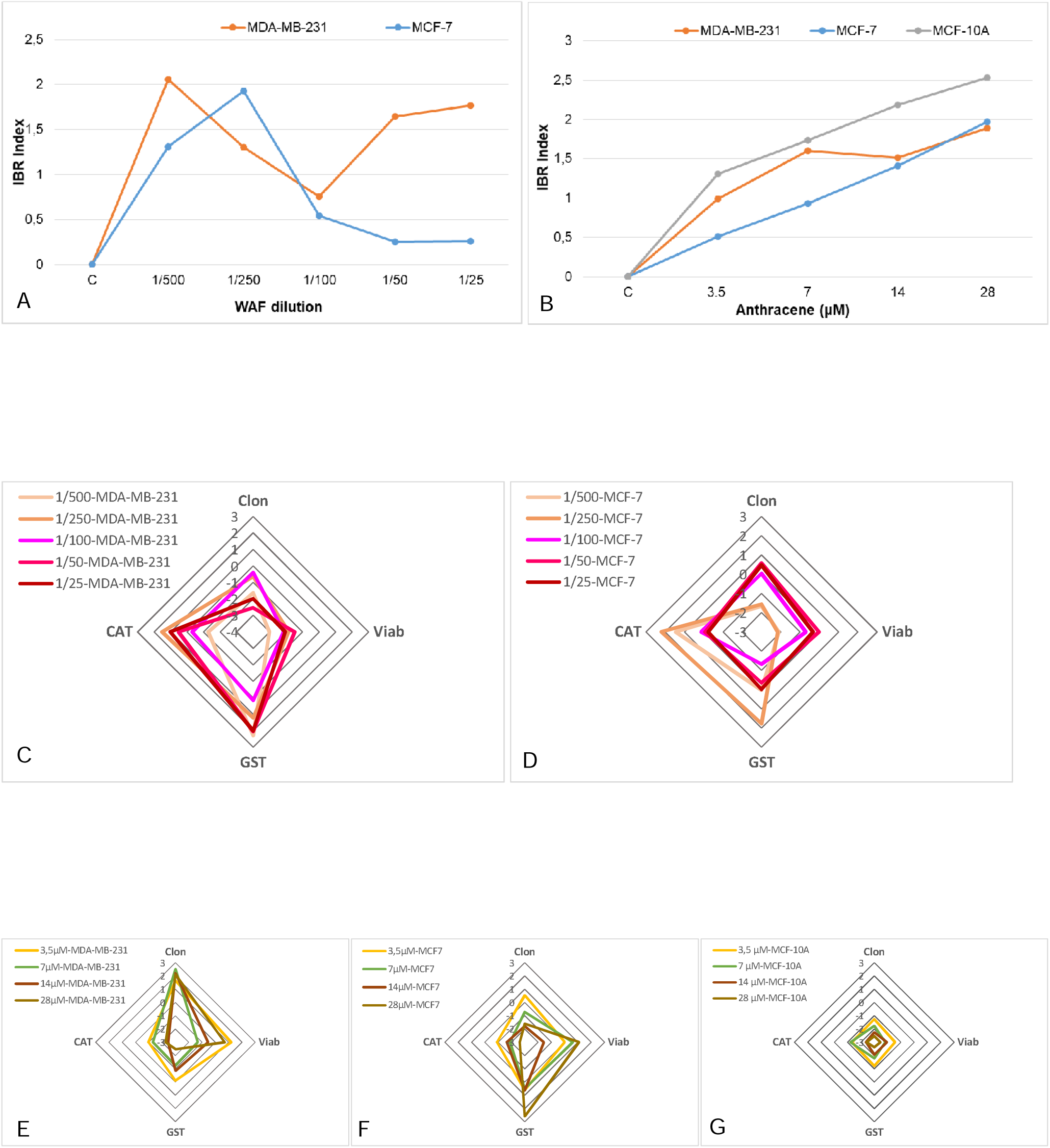
A and B: Integrated biomarker response (IBR) index calculated for each WAF dilution (A) or anthracene concentration (B). C and D: IBR values and associated star plots for MDA-MB-2331 (C) and MCF-7 (D) cells exposed to WAF. E, F and G: IBR values and associated star plots for MDA-MB-2331 (E), MCF-7 (F) and MCF-10A (G) cells exposed to anthracene. CAT: catalase activity, GST: glutathione S-transferase activity, Viab: viability, Clon: clonogenicity.

## DISCUSSION

Following an oil spill, the dispersion of petroleum hydrocarbons in the environment includes the continuous transfer between compartments [4]. Available data on the persistence of hydrocarbons in the environment remains scarce. PAH, a subset of petroleum hydrocarbon components, have environmental half-lives spanning from weeks for lower molecular weight fractions to years for higher molecular weight compounds [4].

Our results on WAF characterization showed that total petroleum hydrocarbons content was 3.66 mg/L, while PAH concentration was 0.016 mg/L, with naphthalene and phenanthrene being the most representative, and toluene as the main BTEX at 0.17 mg/L. As the composition and quantity of hydrocarbons in WAF differs depending on variables like oil type, oil-water ratio, mixing duration, and exposure temperature [31], several authors reported different WAF compositions. On the one hand, del Brio et al., Lavarías et al. and Li et al. reported similar TPH levels in WAF [1,4,32]. Employing identical WAF preparation techniques, del Brio et al. reported a concentration of 2.18_mg/L TPHs in WAF while Lavarías et al. documented a concentration of 3.41_mg/_L for WAF prepared with Punta Loyola (Santa Cruz, Argentina) crude oil [1,4]. Li et al. reported a concentration of 5.23 ± 0.21 mg/L TPH in WAF prepared with crude oil from Oman and 6.47 ± 0.15 μg/L total PAH with naphthalene, phenanthrene, fluorene and acenaphthene being the most abundant [32]. On the other hand, Neff et al. observed concentrations ranging from 0.008 to 38.31_mg/_L across various Australian crude oils[33]. Therefore, it is necessary to study the composition and toxicity of each WAF prepared with different preparation techniques and with petroleum extracted from different regions in order to evaluate the consequences of oil spills in different areas of interest.

In this work we observed that WAF effects are according to the degree of dilution and to the malignant grade of the breast cell line evaluated. Our results show that exposure to the most diluted WAF solutions significantly affected breast cancer cell survival. Although there were no significant differences in MCF-7 clonogenicity, the trend of the response to WAF exposure was similar to that observed for viability, showing the lowest number of colonies in the groups of cells treated with the most diluted WAF solutions. In MDA-MB-231 cells, the response observed was different, showing the lowest rate of clonogenicity for the most diluted (1/500) and for the most concentrated (1/50 and 1/25) WAF solutions. As a response to the redox imbalance caused by WAF exposure, the antioxidant/detoxifying system is activated. In MCF-7 cells, the antioxidant/ detoxifying response was principally triggered by an increase in CAT activity and in MDA-MB-231 cells by an increase in GST activity. However, this response seems not to be enough to manage the toxic effects triggered by WAF at the most diluted solutions in MCF-7 cells and at the most concentrated solutions in MDA-MB-231 cells.

The increase in CAT activity in MCF-7 would help with the detoxification of H_2_O_2_ increased by WAF. In turn, the increment in GST activity would cooperate with the detoxification of the toxic WAF components from the cell. However, our results showing a decrease in MCF-7 viability and MDA-MB-231 viability suggests that the 25% increase in GST activity is not enough to detoxify the toxic components present in WAF mixture leading to an imbalance that could not be compensated. When diluting the mixture, mostly toxic components might remain in the solution. The results on oxidative metabolism could have long-term effects on viability and/or have effects on other pathways that may be transforming/malignant. These oxidative changes can have consequences on macromolecules such as DNA affecting cell survival or transformation.

As WAF is a heterogeneous mixture of hydrocarbons, dilutions can lead to different responses. At the lowest concentrations, compounds that reduce viability may be more present or predominant and those that stimulate proliferation may be less present. On the other hand, at higher concentrations, compounds that damage/oxidize molecules (DNA/proteins/lipids) might be predominant, triggering a decrease in clonogenicity and viability. The response to toxic compounds is not monotonic, since at different concentrations different pathways are activated leading to different responses. The effect ends up being the sum of the concentration and the effect of each of the components present at those concentrations. It is important to highlight that in a complex mixture of hydrocarbon compounds such as WAF, we must consider the effects of the interactions between the compounds to understand and explain the effects on cells, organisms and the environment. Hence the importance of evaluating toxicants in the context of mixtures.

Other authors reported similar effects of WAF on the survival and antioxidant system of several organisms. Lavarías et al. observed significant increases in CAT, SOD, and GST activities in hepatopancreas and CAT activity in gills of the crustacean *Macrobrachium borellii* after 7 days of exposure to a sublethal WAF concentration [34]. Alsaadi et al. reported that CYP1a mRNA levels increased up to 46-fold and GST 3.8-fold in fish after WAF exposure [35]. However, there were no significant changes in the mRNA levels of p53, SOD, CAT and GSR. Hatching rate of the copepod *T. japonicus* was significantly reduced and GST, GR and CAT activities were increased after WAF exposure in a concentration-dependent manner [36]. In the rotifer *Brachionus koreanus*, exposure to WAF resulted in elevated ROS levels, concomitant with reduced lifespan and fecundity [37]. These findings suggest that WAF exposure may induce oxidative stress, leading to adverse effects on reproductive fitness in aquatic organisms. Little is known about the effects of WAF on human cells. In this sense, Major et al. studied the effects of WAF on human airway BEAS-2B epithelial cells. They found more than 70% cell death after exposure to 1000 ppm WAF. At low concentration exposures (100 and 300 ppm) they observed DNA damage [38]. Rodríguez-Trigo studied health changes in fishermen after participation in crude oil clean-up work and they found persistent oxidative stress in the pulmonary system [39]. Ssempebwa et al. found that concentrations of 5, 10, 15, 20, or 25 ppm of new or used oil did not alter the growth of MCF-7 breast cancer cells [40]. Nevertheless, the mechanisms involved in human cells are not fully understood.

After addressing changes in viability and clonogenicity with WAF exposure, we chose anthracene as a representative of PAH. Our results showed a significant decrease in cell viability and clonogenicity only in non-tumorigenic MCF-10A cells in a dose-dependent manner, along with a decrease in GST and CAT activities. When MCF-10A cells were exposed to anthracene solutions higher than 7 μM, the antioxidant system was unable to compensate for the oxidative stress caused by this toxicant, leading to decreased viability and proliferation. On the other hand, anthracene exposure did not significantly affect viability or proliferation in tumorigenic cells MDA-MB-231 and MCF-7 cells, but significant changes were found in CAT and GST activities of both cell lines. In MCF-7 cells, CAT activity decreased after anthracene exposure and therefore it might not adequately detoxify H_2_O_2_, while the increment observed in GST activity suggests that the detoxifying response might be triggered through GST. In MDA-MB-231 cells, although CAT and GST activities were decreased, viability and proliferation were not affected, suggesting that other detoxifying mechanisms could be activated in these cells or other compensation mechanisms might be induced to balance the oxidative situation. Besides, non-tumorigenic MCF-10A cells might be more sensitive to anthracene than tumorigenic MCF-7 and MDA-MB-231 cell lines. Xu et al. described that redox status could be a breast cancer cell marker that may allow us to distinguish between cancer and non-cancer breast tissue [41]. A deeper redox imbalance has been observed in breast cancer tissues compared to normal tissues, which has been explained due to an electron transport activity induced by the reoxygenation of mitochondria [42]. Similar results were observed in cells from other organisms. Sun et al. [43]studied the effects of anthracene-induced oxidative stress and genetic toxicity at molecular and cellular levels in primary coelomocytes of earthworms. Similar to what we showed in this work, they found that exposure to anthracene altered the redox balance in coelomocytes inducing oxidative stress and leading to a reduction in viability and a decrease in mitochondrial membrane potential. Additionally, the comet assay demonstrated DNA strand breaks, revealing anthracene-induced DNA damage [43].

Analyzing the results obtained from the multivariate analysis of responses, we can conclude that non-tumorigenic cells and tumorigenic cells respond differently to the exposure to both contaminants. Furthermore, tumorigenic MCF-7 and MDA-MB-231 cells were differentially affected by the mixture of hydrocarbons present in WAF and by exposure to the single PAH anthracene. Although tumorigenic cells had a similar response in relation to the effect of WAF on viability and proliferation, they presented a differential response of the antioxidant and detoxifying system. MCF-7 cells responded mainly by activating the antioxidant system through CAT when exposed to low concentrations of WAF, while MDA-MB-231 cells also activated the detoxifying system through an increase in GST. Additionally, both cell lines also showed a differential antioxidant/detoxifying response to anthracene exposure: MCF-7 cells responded with high values of GST activity, while MDA-MB-231 decreased the activities of GST and CAT.

Coincidentally with the PCA, the IBR analysis showed for both tumorigenic cell lines a non-monotonic and opposite response to the highest concentrations of WAF. The biphasic response obtained after WAF exposure might be due to the fact that it is a complex mixture of hydrocarbons and that its dilution may differentially affect biomarker levels. This could be because this response depends not only on the concentration of each compound in the mixture, but also on the interaction between them. IBR analysis for anthracene showed a concentration-dependent upward response, for all the cell lines. Notably, for non-tumorigenic cells the IBR values were higher than those for tumorigenic cells, probably related to the fact that anthracene exposure caused a decrease in all the parameters evaluated, as it is uniformly seen in the star plot. The increasing profile of IBR with the concentration, according to the expected behavior of biomarkers, suggests that the combination of cellular viability and antioxidant/ detoxifying enzyme biomarkers represents an adequate battery to determine anthracene and probably other PAH toxicity.

### Concluding remarks

Our results analyzed together allow us to conclude that the sensitivity of cells to the different contaminants evaluated is affected by the degree of cellular tumorigenicity. These findings are especially relevant as it shows that non-transformed cells can be more affected than transformed cells. Nevertheless, transformed cells may also be affected by WAF and/or PAH, aggravating malignant pathological conditions. It is also worth highlighting the benefits of measuring several biochemical and cellular parameters and then analyzing them together through multivariate analysis as biomarkers, obtaining a broader and more comprehensive evaluation of the effects of toxicants.

## Acknowledgments

This work was supported by grants from the Agencia Nacional de Promoción Científica y Tecnológica (ANPCyT) PICT-2019-02355.

Figure Supplementary 1: Principal Component Analysis on biomarker responses assessed in MDA-MB-231, MCF-7 and MCF-10A cells exposed to WAF or anthracene. CAT: catalase activity, GST: glutathione S-transferase activity, Viab: viability, Clon: clonogenicity. MDA: MDA-MB-231 cells, MCF-7: MCF-7 cells. MCF10: MCF-10A cells. WAF dilutions: 1/500, 1/250, 1/100, 1/50, 1/25. Anthracene concentrations: 3.5, 7, 14, 28 µM.

Figure Supplementary 2: Principal Component Analysis on biomarker responses assessed in MDA-MB-231 and MCF-7 cells exposed to WAF or anthracene. CAT: catalase activity, GST: glutathione S-transferase activity, Viab: viability, Clon: clonogenicity. MDA: MDA-MB-231 cells, MCF-7: MCF-7 cells. WAF dilutions: 1/500, 1/250, 1/100, 1/50, 1/25. Anthracene concentrations: 3.5, 7, 14, 28 µM. CW: control WAF, CA: control anthracene.

