## Supplementary figures and images for "Hydrocarbon effects on cell survival and antioxidant system of breast tumorigenic and non-tumorigenic cells"

### Supplementary 2

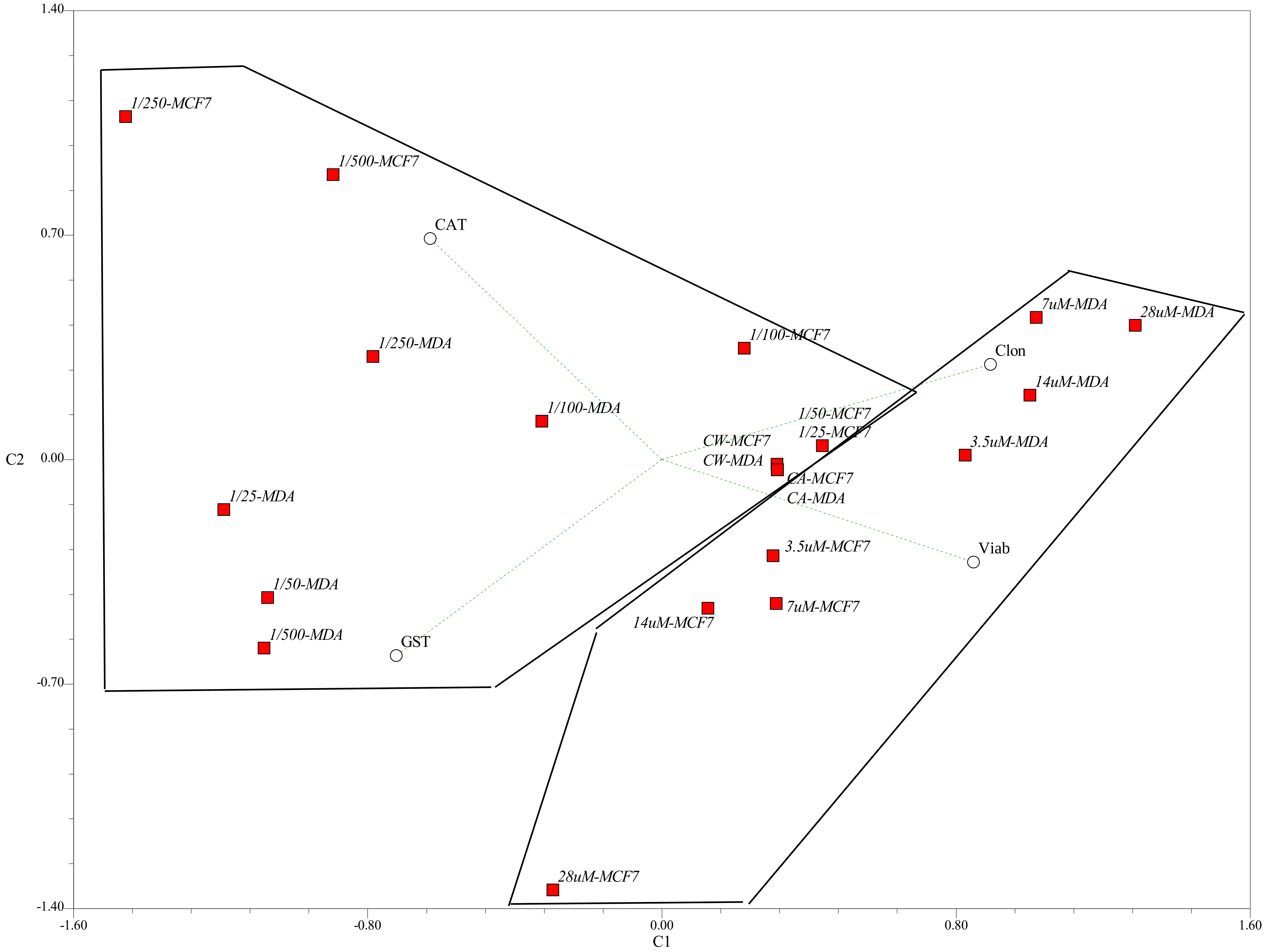

### Supplemetary 1

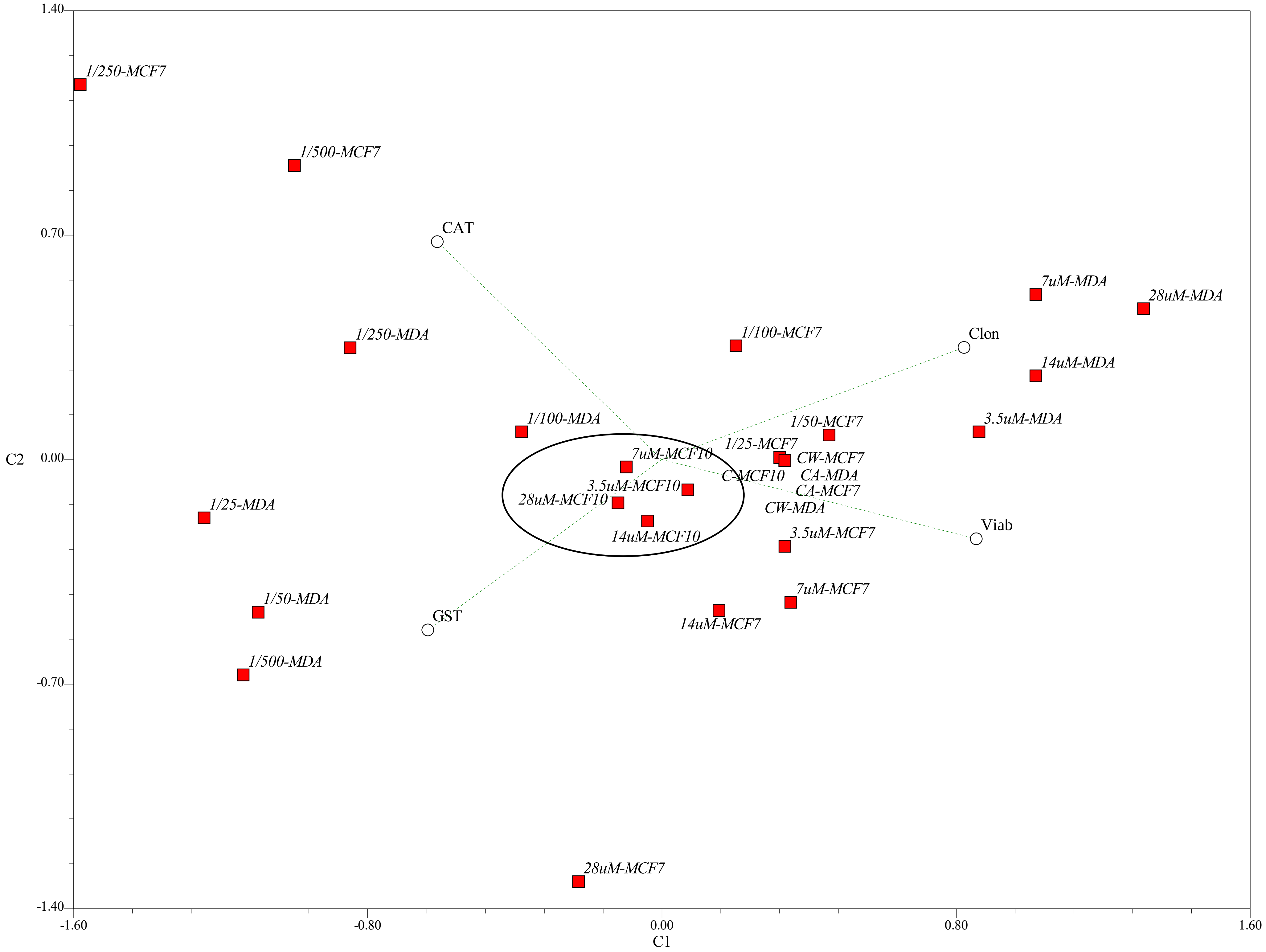
